# Exact distribution of divergence times from fossil ages and tree topologies

**DOI:** 10.1101/490003

**Authors:** Gilles Didier, Michel Laurin

## Abstract

Being given a phylogenetic tree of both extant and extinct taxa in which the fossil ages are the only temporal information (namely, in which divergence times are considered unknown), we provide a method to compute the exact probability distribution of any divergence time of the tree with regard to any speciation (cladogenesis), extinction and fossilization rates under the Fossilized-Birth-Death model.

We use this new method to obtain a probability distribution for the age of Amniota (the synapsid/sauropsid or bird/mammal divergence), one of the most-frequently used dating constraints. Our results suggest an older age (between about 322 and 340 Ma) than has been assumed by most studies that have used this constraint (which typically assumed a best estimate around 310-315 Ma) and provide, for the first time, a method to compute the shape of the probability density for this divergence time.

## INTRODUCTION

Dating the Tree of Life (TOL from here on) has become a major task for systematics because timetrees are increasingly used for a great diversity of analyses, ranging from comparative biology (e.g., Felsenstein 1985, 2012) to conservation biology (Faith 1992), and including deciphering patters of changes in biodiversity through time (e.g., Feng *et al.* 2017). Dating the TOL has traditionally been a paleontological enterprise (Romer 1966), but since the advent of molecular dating (Zuckerkandl and Pauling 1965), it has come to be dominated by molecular biology (e.g., Kumar and Hedges 2011). However, recent developments have highlighted the vital role that paleontological data must continue to play in this enterprise to get reliable results (Heath *et al.* 2014; Zhang *et al.* 2016; Cau 2017; Guindon 2018).

Most of the recent efforts to date the TOL have used the node dating method (e.g., Shen *et al.* 2016), even though several recent studies point out at limitations of this approach (e.g., van Tuinen and Torres 2015), notably because uncertainties about the taxonomic affinities of fossils have rarely been fully incorporated (e.g., Sterli *et al.* 2013), incompleteness of the fossil record is difficult to take into consideration (Strauss and Sadler 1989; Marshall 1994, 2019; Marjanović and Laurin 2008; Nowak *et al.* 2013), or simply because it is more difficult than previously realized to properly incorporate paleontological data into the proper probabilistic framework required for node dating (Warnock *et al.* 2015). More recently, other methods that incorporate more paleontological data have been developed. Among these, tip dating (Pyron 2011; Ronquist *et al.* 2012a,b) has the advantage of using phenotypic data from the fossil record (and molecular data, for the most recent fossils) to estimate the position of fossils in the tree and thus does not require fixing a priori this parameter, and comparisons between tip and node dating are enlightening (e.g., Sharma and Giribet 2014). Even though this method was developed to better date clades, it can improve phylogenetic inference by using stratigraphic data to better place extinct taxa in the phylogeny, as recently shown by Lee and Yates (2018) for crocodylians. Tip dating was first developed in a context of “total evidence” analysis combining molecular, morphological and stratigraphic data, but it can be performed using only paleontological (morphological and stratigraphic) data, as has been done for trilobites (Paterson *et al.* 2019), echinoderms (Wright 2017), theropods (Bapst *et al.* 2016; Lloyd *et al.* 2016), canids (Slater 2015) and hominids (Dembo *et al.* 2015).

Another recently-developed approach is the “fossilized birth-death process” (Stadler 2010; Heath *et al.* 2014), which likewise can incorporate much paleontological data, but using birth-death processes (in addition to the phenotypic or molecular data to place fossils into a tree). It was initially used to estimate cladogenesis, extinction and fossilization rates (Stadler 2010; Didier *et al.* 2012, 2017), but it can also be used to estimate divergence times (Heath *et al.* 2014). The approach of Heath et al. (2014) differs from ours in how it incorporates fossils into the analysis and in using a Bayesian approach with MCMC to estimate parameter values and divergence times, rather than the exact computations that we present. Both differences deserve comments.

Heath *et al.* (2014) chose to incorporate only part of the phylogenetic information available for some fossils. Namely, for each fossil, the node that it calibrates must be identified. That “ calibration node” is the most recent node that includes the hypothetical ancestor of the fossil in question. This approach has both advantages and disadvantages compared with our approach. The advantage is that this method can incorporate fossils whose placement in the Tree of Life is only approximately known, and character data about extinct taxa need not be collected for this approach. The drawbacks include the following: this approach prevents dating fully-extinct clades (something that our method can handle), and in the most-intensively studied clade, this procedure may discard useful information. That method introduces a different procedure to handle extinct vs. extant taxa, whereas both are part of the same Tree of Life. Indeed, the approach of Heath *et al.* (2014) seems to take for granted that the phylogenetic position of extinct taxa is inherently more poorly documented than that of extant taxa, but this may not always be the case. For instance, at the level of the phylogeny of tetrapods, the position of lissamphibians is more controversial than that of the many of their presumed Paleozoic relatives (Marjanović and Laurin 2019). And the long branches created by the extinction of intermediate forms (visible only in the fossil record, when preserved) can also create problems in phylogenetic inference of extant taxa (Gauthier *et al.* 1988). Our approach uses a population of trees to represent phylogenetic uncertainty of extant and extinct taxa simultaneously. Consequently, it requires either character data for extinct and extant taxa, or a population of trees for these taxa obtained from the literature.

Heath *et al.* (2014) use Monte Carlo Markov Chain (MCMC) to estimate parameter values and divergence times, but we present (below) a method to calculate exactly the distribution of divergence times from the diversification and fossilization rates. Note that our computation can be easily integrated in a MCMC scheme like that of Heath *et al.* (2014). The resulting approach would obtain results more accurate in a shorter time since it does not have to sample the divergence times but only the rates, the tree topology and the fossils placement. In the present work, we devised a MCMC-based importance sampling procedure to deal with the uncertainty on the model rates and on the fossil ages (Appendix D).

Another method that uses birth and death models is cal3 (Bapst 2013; Bapst and Hopkins 2017), which has been developed specifically to time-scale paleontological trees. It differs from our approach in at least three important points. The first concerns the limits of the validity of the method; as we explained in Didier *et al.* (2017), cal3 assumes that the diversification process extends over unlimited time, and it approximates the correct solution only in extinct taxa. By contrast, our method can handle trees that combine extant and extinct taxa. The second difference is that cal3 can test for ancestral position of some taxa, though it uses only the temporal range of taxa for this, disregarding character data (Bapst and Hopkins 2017:64). By contrast, we did not yet implement an algorithm to check for ancestors preserved in the fossil record, but we did this manually by considering stratigraphic and character data (Didier *et al.* 2017); we consider, in a first approximation, that an ancestor should be geologically older than its descendants and that it should not display a derived character that is absent from its putative descendant. The third important difference is that cal3 does not incorporate a method to estimate the birth-and-death model parameters (cladogenesis, extinction and sampling rates). Instead, it relies on previously-published estimates of these rates, or these can be obtained through other methods. For instance, Bapst and Hopkins (2017) obtained their estimates using paleotree (Bapst 2012), which implements a method proposed by Foote (1997). That method uses stratigraphic ranges of taxa in time bins to infer rate parameters. Otherwise, such estimates in the literature are relatively rare, and most published rates were published as “per capita” origination and extinction rates using the method developed by Foote (2000). That method is a taxic approach that does not require a phylogeny and that quantifies the turnover rate between taxa at a given rank. Given that some taxa may be paraphyletic, and that the taxonomic (Linnaean) ranks are subjective (Ereshefsky 2002; Laurin 2008), converting these rates into cladogenesis, extinction and fossilization (sampling) rates is not straightforward. Indeed, Bapst (2014: 346) acknowledged these problems and concluded that “These rate estimates cannot always be obtained, particularly in fossil records consisting entirely of taxonomic point occurrences, such as many vertebrate fossil records. Further advances in time-scaling methods are needed to deal with these datasets”. Below, we present such advances.

Our method relies on an assessment of these rates based on detailed phylogenetic and stratigraphic data (Didier *et al.* 2017). Instead of stratigraphic ranges and time bins, our method requires more detailed stratigraphic information about the age of every horizon that has yielded fossils of each taxon included in the analysis, or a random sample thereof. This requires greater data collection effort and may be limiting in some contexts, but the more detailed data should improve the accuracy of the method. This is supported by the simulations performed by (Foote 1997), which showed that preservation rate and completeness are better estimated (with less variance around the true value) when all stratigraphic occurrences are used, rather than only ranges; similarly, the simulations of (Heath *et al.* 2014: fig. 2B) shows that using all fossils (rather than only the oldest ones of each clade) reduces the confidence intervals of estimated node ages. Thus, our method can take full advantage of the many detailed paleontological phylogenies that have been published recently and recent progress in geochronology, but it cannot be applied for taxa that lack such a well-studied fossil record of phenotypically-complex organisms that can be placed fairly precisely in a phylogeny and in the stratigraphy.

This contribution develops a new method to date the TOL using the fossilized birth-death process. This method relies on estimating parameters of the fossilized birth-death process through exact computations and using these data to estimate a probability distribution of node ages. This method currently requires input trees, though it could be developed to include estimation of these trees. These trees include fossils placed on terminal or on internal branches, and each occurrence of each taxon in the fossil record must be dated. In this implementation, we use a flat probability distribution of fossil ages between two bounds, but other schemes could easily be implemented. Our new method thus shares many similarities with what we recently presented (Didier *et al.* 2017), but it is aimed at estimating nodal ages rather than rates of cladogenesis, extinction, and fossilization. Namely, in Didier *et al.* (2017), we proposed a method for computing the probability density of a phylogenetic tree of extant and extinct taxa in which the fossil ages was the only temporal information. The present work extends this approach in order to provide an exact computation of the probability density of any divergence times of the tree from the same information.

We then use these new developments to estimate the age of the divergence between synapsids and sauropsids, which is the age of Amniota, epitomized by the chicken/human divergence that has been used very frequently as a dating constraint (e.g., Hedges *et al.* 1996; Hugall *et al.* 2007). In fact, Müller and Reisz (2005: 1069) even stated that “The origin of amniotes, usually expressed as the ‘mammal-bird split’, has recently become either the only date used for calibration, or the source for ‘secondary’ calibration dates inferred from it.” The recent tendency has fortunately been towards using a greater diversity of dating constraints, but the age of Amniota has remained a popular constraint in more recent studies (e.g., Hugall *et al.* 2007; Marjanović and Laurin 2007; Shen *et al.* 2016). However, Müller and Reisz (2005: 1074) argued that the maximal age of Amniota was poorly constrained by the fossil record given the paucity of older, closely related taxa, and that this rendered its use in molecular dating problematic. Thus, we believe that a more sophisticated estimate of the age of this clade, as well as its probability distribution, will be useful to systematists, at least if our estimates provide a reasonably narrow distribution. Such data are timely because molecular dating software can incorporate detailed data about the probability distribution (both its kind and its parameters) of age constraints (Ho and Phillips 2009; Sauquet 2013), the impact of these settings on the resulting molecular age estimates is known to be important (e.g., Warnock *et al.* 2012), but few such data are typically available, though some progress has been made in this direction recently (e.g., Nowak *et al.* 2013).

The approach presented here was implemented as a computer program and as a R package, which both the probability distribution of divergence times from fossil ages and topologies and are available at https://github.com/gilles-didier/DateFBD.

## METHODS

### Birth-death-fossil-sampling model

We consider here the model introduced in Stadler (2010) and referred to as the Fossilized-Birth-Death (FBD) model in Heath *et al.* (2014). Namely, speciations (cladogeneses) and extinctions are modelled as a birth-death process of constant rates *λ* and *μ*. Fossilization is modelled as a Poisson process of constant rate *ψ* running on the whole tree resulting from the speciation-extinction process. In other words, each lineage alive at time *t* leaves a fossil dated at *t* with rate *ψ*. Last, each lineage alive at the present time, i.e., each extant taxon, is sampled with probability *ρ*, independently of any other event. In practice, “sampled” may mean discovered or integrated to the study. Note that, in constrast with our previous works (Didier *et al.* 2012, 2017) which were based on the FBD model with full sampling of extant taxa (i.e., with *ρ* = 1), we shall consider uniform sampling of the extant taxa in the present work. In all what follows, we make the usual technical assumptions that *λ*, *μ*, *ψ* and *ρ* are non-negative, that *λ* > *μ*, and that *ψ* and *ρ* are not both null. Unlike the FBD process as it is sometimes described, we consider the usual “from past to present” time direction in this work.

We shall not deal directly with the whole realizations of the FBD process (Figure 1-left) but rather, with the part of realizations which can be reconstructed from present time using data from extant taxa and from the fossil record; this will be referred to as the *(reconstructed) phylogenetic tree with fossils*. Indeed, we have no information about the parts of the diversification process leading to extinct taxa that left no fossil record or non-sampled extant or extinct species (Figure 1-center). Let us state more precisely what is assumed to be reconstructible. We make the assumption that the starting time of the diversification (i.e., the base of the root branch) and the fossil ages are known (in practice, we specify these as intervals, over which we sample randomly using a flat distribution for fossil ages, and we integrate for the root age) and that we are able to accurately determine the phylogenetic relationships between all the extant and extinct taxa considered, in other words, that we can reconstruct the actual tree topology of the species phylogeny (Figure 1-right). Note that under these assumptions, the only available temporal information in a reconstructed phylogenetic tree with fossils is given by the fossil ages, the starting time of the diversification process (which is the only temporal information that is poorly constrained) and the present time. Namely, all the divergence times of the reconstructed phylogeny are unknown.

**Figure 1:**
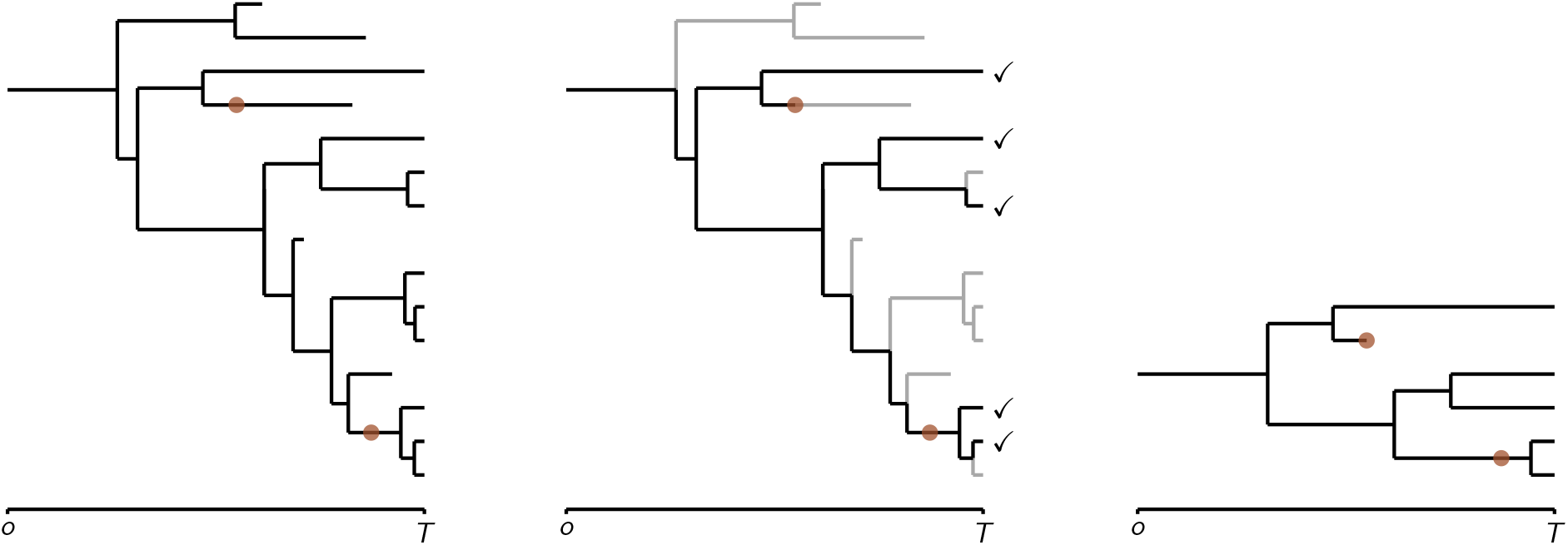
Left: a realization of the diversification process with fossil finds (represented by “•”); Center: its observable part from the sampled extant taxa and the fossil finds is displayed in black – the gray parts are lost (sampled extant species are those with “✓”); Right: the resulting reconstructed phylogenetic tree.

**Figure 2:**
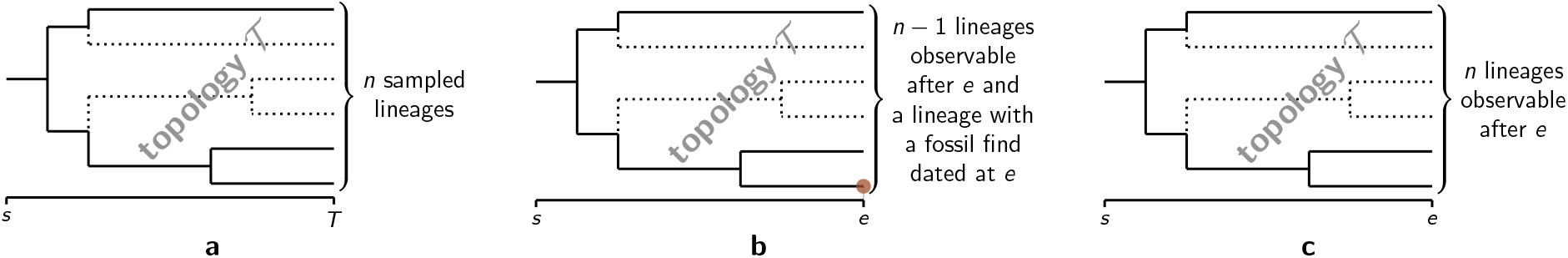
The three types of patterns on which our computations are based. The patterns of type **a** end at the present time *T* while the patterns of type **b** and **c** are parts of the diversification process that continue after *e* < *T*. Patterns of type **c** end with the period covered by a study (before the present). Here, “observable” stands for “observable at *e*”. Note that the bottom-most lineage in type **b** may or may not be observable (strictly) after *e*.

Most of our calculations are based on probabilities of the following basic events, already derived in (Stadler 2010; Didier *et al.* 2012, 2017) under the FBD model.

The probability that a single lineage starting at time 0 has *n* descendants sampled with probability *ρ* at time *t* > 0 without leaving any fossil (i.e., neither from itself nor from any of its descendants) dated between 0 and *t* is given by

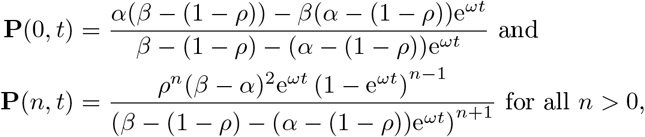

where *α* < *β* are the roots of −*λx*^2^ + (*λ* + *μ* +*ψ*)*x* − *μ* = 0, which are always real (if *λ* is positive) and are equal to

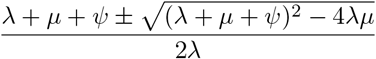

and *ω* = −*λ*(*β α*).

We showed in Didier *et al.* (2017) that *α* can be interpreted as the asymptotical proportion of lineages unobservable from the fossil record, assuming the the diversification process continues indefinitely. This quantity *α* is close to the complementary of the probability *P_s_* defined in Bapst (2013) and used in Wagner (2019). We did not find biological interpretations of the quantities *ω* and *β*, which are essentially convenient to write the computations below. A table of the notations is provided in Appendix A.

Let us write **P_o_**(*t*) for the probability that a lineage present at time 0 is observable at *t*, which is the complementary probability that it both has no descendant sampled at the present time *T* and lacks any fossil find dated after *t*. Namely, we have that

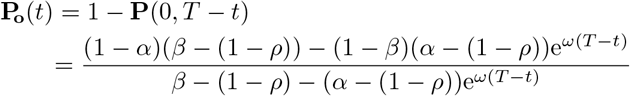

### Tree topologies

We shall consider only binary, rooted tree topologies since so are those resulting from the FBD process. Moreover, all the tree topologies considered below are *labelled* in the sense that all their tips are unambiguously identified. From now on, “tree topology” will refer to “labelled-binary-rooted tree topology”.

Let us use the same notations as Didier (2020). For all tree topologies 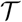, we still write 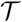 for the set nodes of 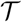. We put 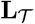 for the set of tips of 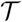 and, for all nodes *n* of 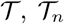 for the subtree of 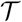 rooted at *n*. For all sets *S*, |*S*| denotes the cardinality of *S*. In particular, 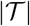 denotes the size of the tree topology 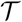 (i.e., its total number of nodes, internal or tips) and 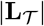 its number of tips.

#### Probability

Let us define 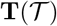 as the probability of a tree topology 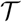 given its number of tips under a lineage-homogeneous process with no extinction, such as the reconstructed birth-death-sampling process.

##### Theorem 1

(Harding 1971). *Given its number of tips, a tree topology* 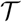 *resulting from a pure-birth realization of a lineage-homogeneous process has probability* 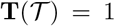 *if* 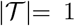, *i.e.*, 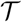 *is a single lineage. Otherwise, by putting a and b for the two direct descendants of the root of* 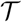 *, we have that*

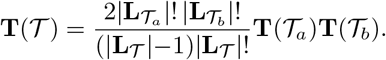

The probability provided in Didier *et al.* (2017, Supp. Mat., Appendix 2) is actually the same as that just above though it was derived in a different way from Harding (1971) and expressed in a slightly different form (Appendix B).

From Theorem 1, 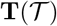 can be computed in linear time through a post-order traversal of the tree topology 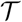.

### Patterns

As in Didier *et al.* (2017), our computations are based on three types of subparts of the FBD process which all start with a single lineage and end with three different configurations, namely

- patterns of type **a** end at the present time *T* with *n* ≥ 0 sampled lineages,
- patterns of type **b** end at a time *e < T* with a fossil find dated at *e* and *n* − 1 ≥ 0 lineages observable at *e*,
- patterns of type **c** end at a time *e < T* with *n* > 0 lineages observable at *e* (the case where *n* = 0 is not required in the calculus).

The probability density of a pattern of any type is obtained by multiplying the probability density of its ending configuration, which includes its number of tips, by the probability of its tree topology given its number of tips provided by Theorem 1.

In this section, we provide equations giving the probability density of ending configurations of patterns of types **a**, **b** and **c**, first with regard to a given “punctual” starting time and, second, by integrating this probability density uniformly over an interval of possible starting times in order to take into account the uncertainty associated with the timing of the beginning of the diversification process.

#### *Patterns of type* **a**

The probability density of the ending configuration of a pattern of type **a** is that of observing *n* ≥ 0 lineages sampled at time *T* by starting with a single lineage at time *s* without observing any fossil between *s* and *T*. We have that

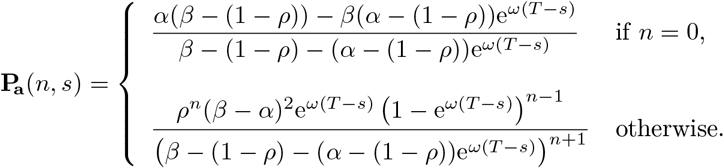

#### *Patterns of type* **b**

The probability density of the ending configuration of a pattern of type **b** is that of observing a fossil find dated at *e* with *n* − 1 ≥ 0 other lineages observable at time *e* by starting with a single lineage at time *s* without observing any fossil between *s* and *e*. From Didier *et al.* (2017), we have that

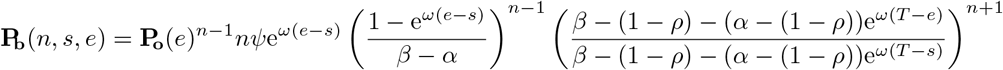

#### *Patterns of type* **c**

The probability density of the ending configuration of a pattern of type **c** is that of getting *n* > 0 lineages observable at time *e* by starting with a single lineage at time *s* without observing any fossil between *s* and *e*. From Didier *et al.* (2017), we have that

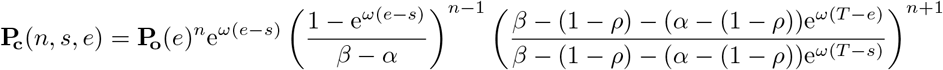

### Basic trees

Following Didier *et al.* (2017), we split a reconstructed realization of the FBD process, i.e. a phylogenetic tree with fossils, by cutting the corresponding phylogenetic tree at each fossil find. The trees resulting of this splitting will be referred to as *basic trees* (Figure 3). If there are *k* fossils, this decomposition yields *k* + 1 basic trees. By construction, basic trees (i) start with a single lineage either at the beginning of the diversification process or at a fossil age, (ii) contain no internal fossil finds and (iii) are such that all their tip-branches either terminate with a fossil find or at the present time. Note that a basic tree may be unobservable (Figure 3). Since tips of basic trees are either fossil finds or extant taxa, they are unambiguously labelled. The set of basic trees of a phylogenetic tree with fossils is a partition of its phylogenetic tree in the sense that all its nodes belong to one and only one basic tree.

**Figure 3:**
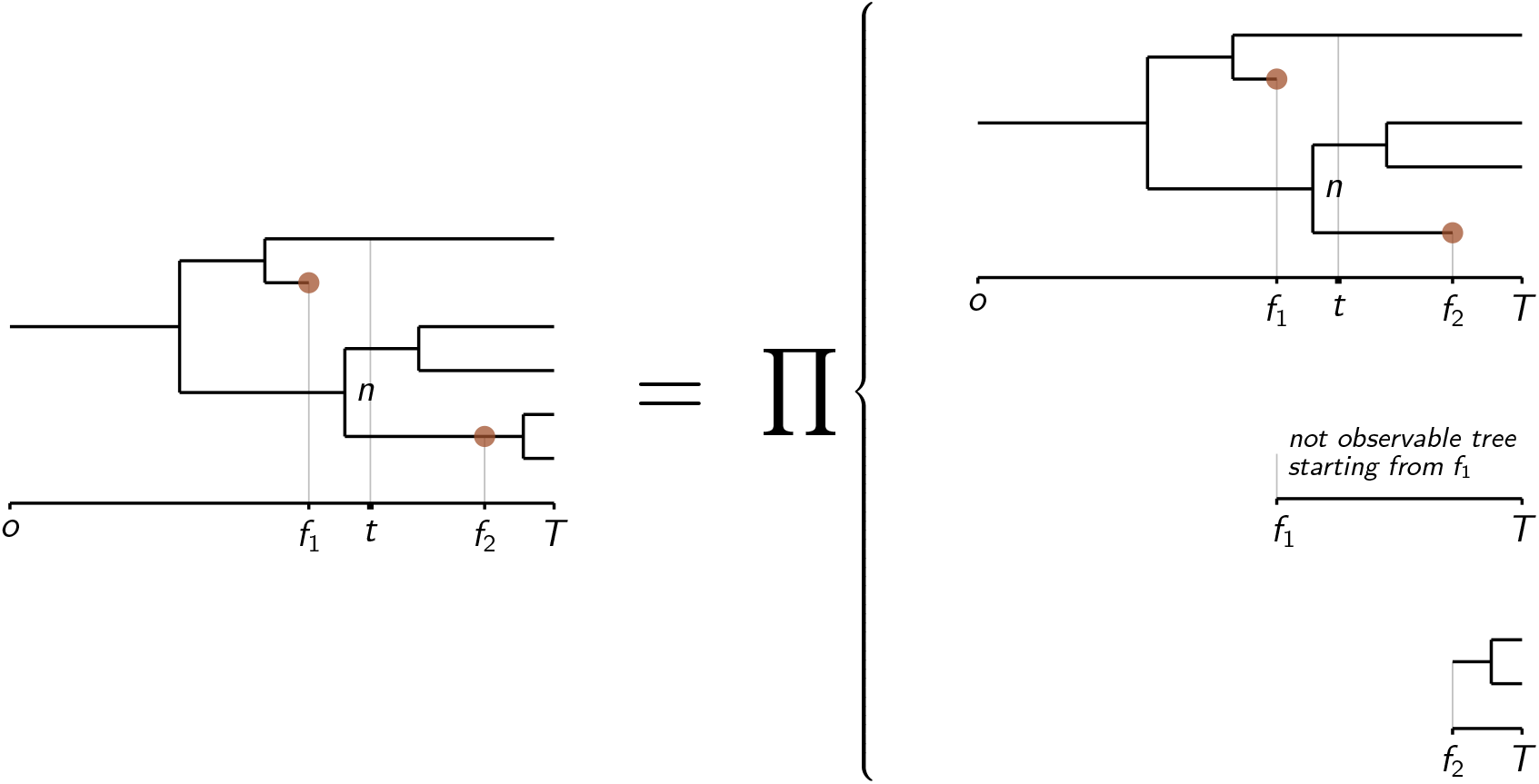
Decomposition of the reconstructed tree of Figure 1 into basic trees, obtained by cutting it at all fossils finds. Remark that the second basic tree from the top in the last column, which starts just after the fossil find dated at *f*_1_, is not observable.

**Figure 4:**
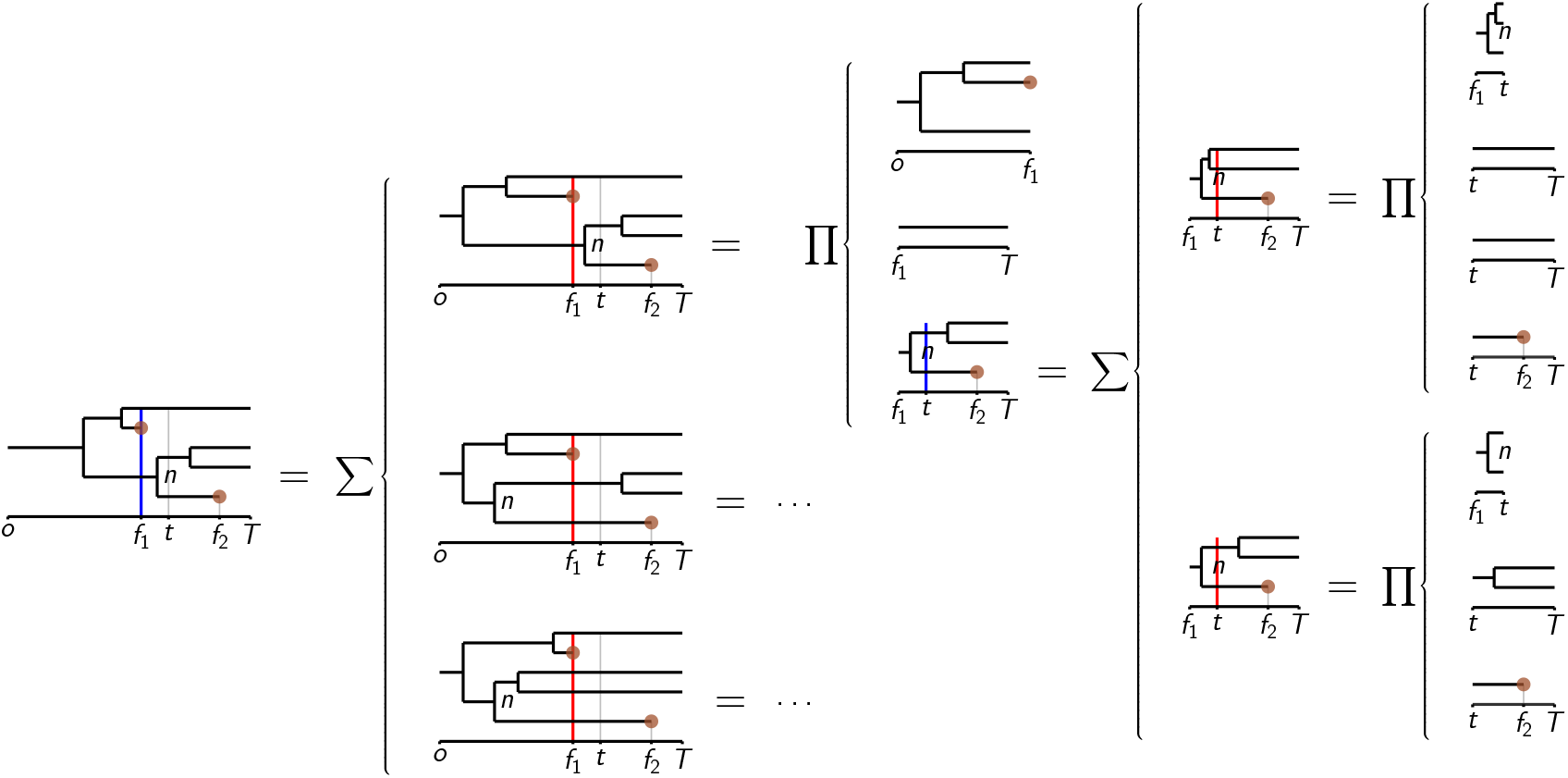
Schematic of the computation of the probability density of the basic tree in the left-most column with the time constraint *τ_n_* ≤ *t* in the case where *t* is posterior to the oldest fossil age *f*_1_. If a given time is displayed in blue, the positions of the nodes of the basic trees delimited by that time say nothing about the position of the corresponding divergences times relative to this time. Conversely, if a given time is displayed in red, all nodes to the left (resp. to the right) of this time have divergence anterior (resp. posterior) to that time. Column 2 displays the three possible sets of anterior nodes consistent with this basic tree and the fossil find dated at *f*_1_. Column 3 shows the computation of the probability density associated to the top one (the two other computations are similar), as a product of probability densities of a pattern of type **b**, a pattern of type **a** and of a basic tree. Column 4 schematizes the computation of this basic tree (Column 3, Row 3), which requires to consider the two possible sets of anterior nodes with regard to the time constraint *τ_n_* ≤ *t* displayed at Column 4 and eventually expressed as products of patterns of types **c**, **a** and **b** (Column 5).

The interest of the decomposition into basic trees stands in the fact that a fossil find dated at a time *t* ensures that the fossilized lineage was present and alive at *t*. Since the FBD process is Markov, the evolution of this lineage posterior to *t* is independent of the rest of the evolution, conditionally on the fact that this lineage was present at *t*. It follows that the probability density of a reconstructed realization of the FBD process is the product of the probability densities of all its basic trees (Didier *et al.* 2017).

Remark that a basic tree is fully represented by a 3-tuple 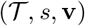, where 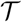 is its topology, *s* is its starting time and **v** is the vector of its tip ages. Namely, for all 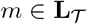, **v**_*m*_ is the age of *m*. We assume that all fossil ages are strictly anterior to the present time. Under the FBD model, the probability that two fossils are dated at the exact same time is zero. In other words, if for two leaves *m* and *m*′ with *m* ≠ *m*′, we have both **v**_*m*_ < *T* and **v**_*m′*_ < *T* then **v**_*m*_ ≠ **v**_*m′*_. For all subsets 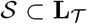, we put 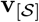 for the vector made of the entries of **v** corresponding to the elements of 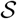.

### Probability distribution of divergence times

Let us consider a phylogenetic tree with fossils 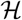 in which the only temporal information is the diversification starting time and the fossil ages, and a possibly empty set of time constraints 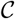 given as pairs {(*n*_1_, *t*_1_), …, (*n*_*ℓ*_, *t*_*ℓ*_)} where *n*_1_, …, *n*_*ℓ*_ are internal nodes of the phylogenetic tree (each one occurring in a single pair of the set) and *t*_1_, …, *t*_2_ are times. For all internal nodes *n* of 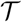, we put *τ_n_* for the divergence time associated to *n*. For any subset of nodes 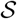, we write 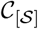 for the set of the time constraints of 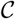 involving nodes in 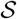, namely 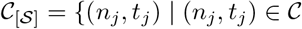 and 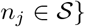}.

We shall see here how to compute the joint probability density of 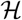 and the events 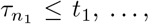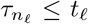, denoted 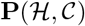. Computing the joint probability of 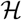 and events 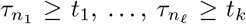 is symmetrical.

Note that the probability distribution of the divergence time associated to a node *n* at any time *t* is given as the ratio between the joint probability density of 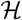 and the event *τ_n_* ≤ *t* to the probability density of 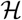 (with no time constraint). Though calculating the distribution of the divergence time of *n* requires only the probability densities of 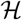 and that of 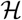 and *τ_n_* ≤ *t* (i.e., 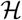 without time constraint and with a single time constraint), we present here the more general computation of the joint probability density of a phylogenetic tree with fossils and an arbitrary number of time constraints since it is not significantly more complicated to write.

From the same argument as in the section above, 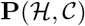, the joint probability density of 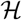 and the events 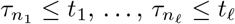 is the product of the probability densities of the basic trees resulting from the decomposition of 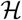 with the corresponding time constraints.

Namely, by assuming that there are *k* fossils in 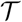 and by putting 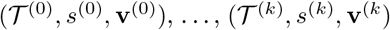 for the basic trees of 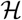 we have that

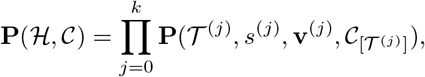

where 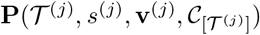 is the probability density of 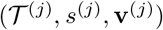 with root or nodal time constraints 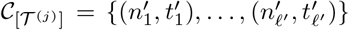, in other words, the joint probability density of the basic tree 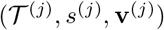 and the events 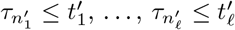.

In order to compute the probability density of a basic tree 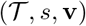 with a set of time constraints 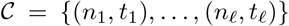, let us define its *oldest age z* as *z* = min{min_*i*_ **v**_*i*_, min_*j*_ *t*_*j*_} and its set of *anterior nodes X* as the union of the nodes *n_j_* such that 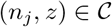 and, if there exists a tip *c* such that **v**_*c*_ = *z*, of the direct ancestor of *c* (under the FBD model, if such a tip exists, it is almost surely unique).

Stating our main theorem requires some additional definitions. Following Didier (2020), a *start-set* of a tree topology 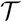 is a possibly empty subset *A* of internal nodes of 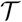 which verifies that if an internal node of 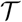 belongs to *A*, then so do all its ancestors.

Being given a tree topology 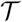 and a non-empty start-set *A*, the *start-tree* 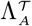 is defined as the subtree topology of 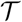 made of all nodes in *A* and their direct descendants. By convention, 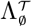, the start-tree associated to the empty start-set, is the subtree topology made only of the root of 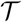.

For all internal nodes *n* of the tree topology 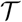, we define 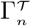 as the set of all the start-sets of 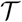 that contain *n*.

#### Theorem 2.

*Let* 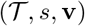 *be a basic tree, T the present time*, 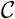 *a possibly empty set of time constraints, z the corresponding oldest age and X the set of anterior nodes. By setting* 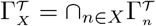, *the probability density of the basic tree* 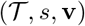 *with time constraints* 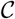 *is*

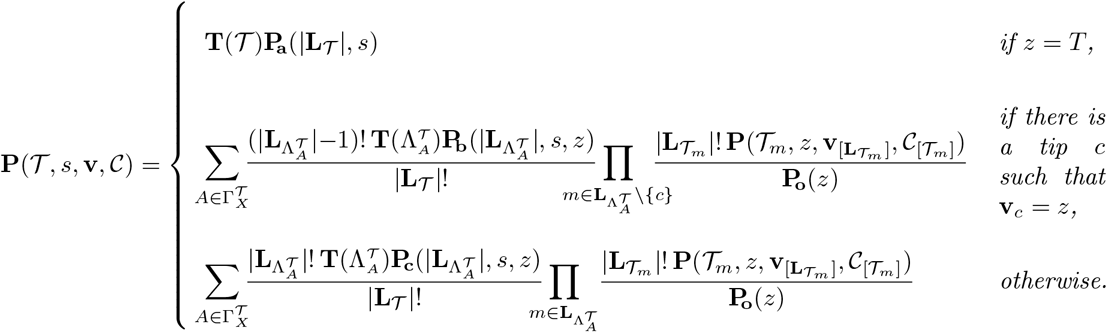

*Proof.* Let us first remark that if *z* = *T*, then we have that 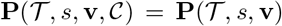, since even if 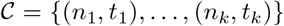 is not empty, *z* = *T* implies that all the times *t_j_* are posterior or equal to *T*, thus we have always 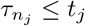. Moreover, the fact that *z* = *T* implies that the basic tree 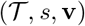 contains no fossil. By construction, it is thus a pattern of type **a** and its probability density is 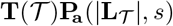.

Let now assume that *z < T* (i.e., the subtree includes a fossil or a time constraint) and that there is a tip *c* associated to a fossil find dated at *z* (i.e., such that **v**_*c*_ = *z*). Let us write the probability density 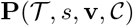 as the product of the probability densities of the part of the diversification process which occurs before *z* and of that which occurs after *z*. Delineating these two parts requires to determine the relative time positions of all the divergences with regard to *z*. Let us remark that some of the divergence times are constrained by the given of the problem. In particular, all the divergence times of the ancestral nodes of *c* are necessarily anterior to *z* and so are the divergence times of the nodes *n_j_* (and their ancestors) involved in a time-constraint such that *t_j_* = *z*. Conversely, the divergence times of all the other nodes may be anterior or posterior to *z*. We thus have to consider all the sets of nodes anterior to *z* consistent with the basic tree and its time constraints. From the definition of the set *X* of anterior nodes associated to 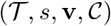, the set of all these sets of nodes is exactly 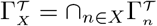. Since all these sets of nodes correspond to mutually exclusive possibilities, the probability density 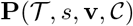 is the sum of the probability densities associated to all of them.

Let us first assume that there exists a fossil find dated at *z*. Given any set of nodes 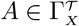 anterior to *z*, the part of diversification anterior to *z* is then the pattern of type **b** starting at *s* and ending at *z* with topology 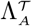, while the part posterior to *z* is the set of basic trees starting from time *z* inside the branches bearing the tips of 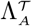 except *c*, i.e, 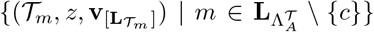, with the corresponding time constraints derived from 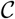, i.e., 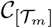 for all 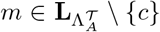 (Figure 5). From the Markov property, the diversification occurring after *z* of all lineages crossing *z* is independent of any other events conditionally on the fact that this lineage was alive at *z*. The probability density of the corresponding basic trees has to be conditioned on the fact that their starting lineage are observable at *z*, i.e., it is 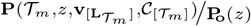 for all nodes 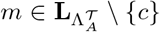. The probability density of the ending configuration of the pattern of type 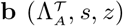 is 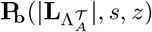. We have to be careful while computing the probability of the topology 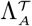 since its tips except *c* (the one associated to the oldest fossil find) are not directly labelled but only known with regard to the labels of their tip descendants while Theorem 1 provides the probability of a (exactly) labelled topology. In order to get the probability of 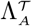, we multiply the probability of 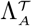 assuming that all its tips are labelled with the number of ways of connecting the tips of 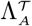 except the one with the fossil (i.e., 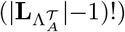) to its pending lineages and the probability of their “labelling”. Since assuming that all the possible labellings of 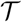 are equiprobable, the probability of the “labelling” of the lineages pending from 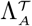 is

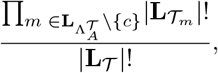

the probability of the tree topology 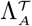 is eventually

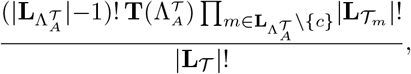

i.e., the product of the probability of the labelled topology 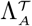 with the number of ways of connecting the (not fossil) tips to the pending lineages and the probability of their “labelling”.

**Figure 5:**
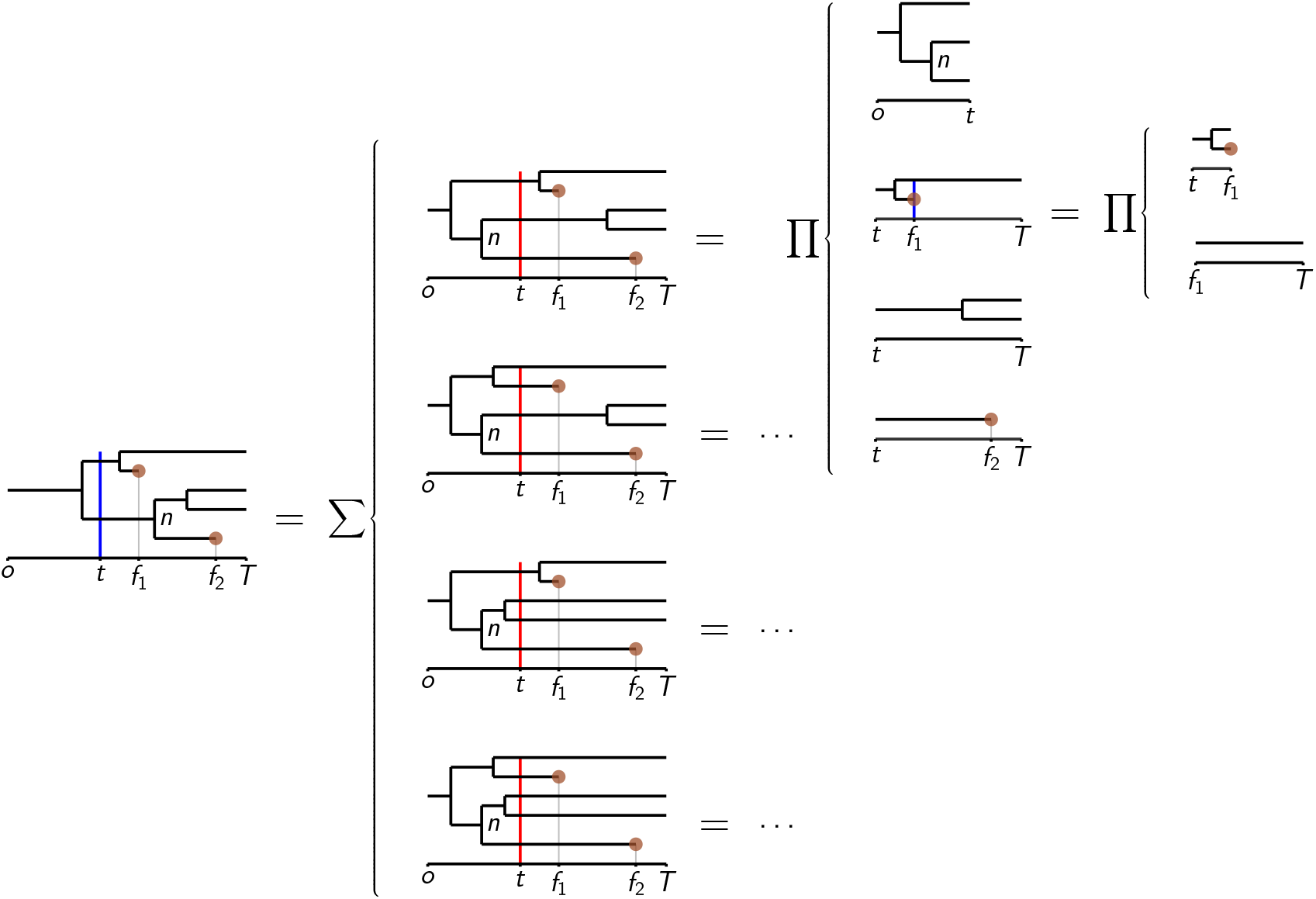
Schematic of the computation of the probability density of the basic tree in the left-most column with the time constraint *τ_n_* ≤ *t* in the case where *t* is anterior to the oldest fossil age *f*_1_. If a given time is displayed in blue, the positions of the nodes of the basic trees delimited by that time say nothing about the position of the corresponding divergences times relative to this time. Conversely, if a given time is displayed in red, all nodes to the left (resp. to the right) of this time have divergence anterior (resp. posterior) to that time. Column 2 displays the four possible sets of anterior nodes consistent with this basic tree and the time constraint *τ_n_* ≤ *t* (i.e., all the possibilities such that the divergence time of node *n* is anterior to *t*). Column 3 shows the computation of the probability density associated to the top one. Column 4 displays the computation of the basic tree at Column 3, Row 2 (the other rows of Column 3 are patterns of type **a**.

The case where *z < T* and where no fossil is dated at *z* is similar. It differs in the fact that the diversification occurring before *z* is a pattern of type **c** (Figure 5) and that the probability of 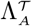 is in this case

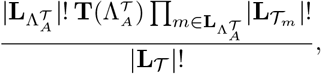

i.e., the product of the probability of the labelled topology 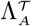 with the number of ways of connecting its tips to the pending lineages (i.e., 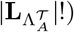) and the probability of their “labelling” (i.e, 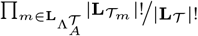).

Theorem 2 allows us to express the probability density of a basic tree with a set of time constraints as a sum-product of probability densities of patterns of type **a**, **b** or **c** and of smaller basic trees, which can themselves be expressed in the same way. Since the basic trees involved in the right part of the equation of Theorem 2 contain either at least one fewer time constraint and/or one fewer fossil find than the one at the left part, this recursive computation eventually ends. We shall not discuss algorithmic complexity issue here but, though the number of possibilities “anterior/posterior to the oldest age” to consider can be exponential, the computation can be factorized in the same way as in Didier (2020) in order to get a polynomial, namely cubic, algorithm.

In the case where the dataset is limited to a period which does not encompass the present (as for the dataset below), some of the lineages may be known to be observable at the end time of the period but data about their fate after this time may not have been entered into the database. As in Didier *et al.* (2017, Section “Missing Data”), adapting the computation to this case is done by changing the type of all the patterns of type **a** to type **c**.

Note that applying the computation with an empty set of time constraints yields to the probability density of a phylogenetic tree with fossils (without the divergence times) as provided by Didier *et al.* (2017). On top of improving the computational complexity of the calculus, which was exponential in the worst case with Didier *et al.* (2017), the method provided here corrects a mistake in the calculus of Didier *et al.* (2017), which did not take into account the question of the labelling and the “rewiring” of the tips of internal basic trees, missing correcting factors in the sum-product giving the probability density that are provided here. Fortunately, this has little influence on the accuracy of rate estimation (Appendix C). In particular, rates estimated from the Eupelicosauria dataset of Didier *et al.* (2017) are less than 5% lower, thus essentially the same, with the corrected method.

For all phylogenetic trees with fossils 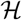, the distribution *F_n_* of the divergence time corresponding to the node *n* of 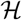 is defined for all times *t* by

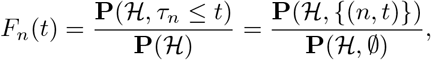

i.e., the probability density of observing 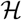 and the divergence of *n* posterior to *t* divided by the probability density of 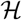 with no constraint. It follows that *F_n_*(*t*) can be obtained by applying the recursive computation derived from Theorem 2 on 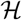 twice: one with the time constraint (*n, t*) and one without time constraints.

### Dealing with time and rate uncertainty

The computation presented in the previous section requires all the fossil ages and the origin of the diversification to be provided as punctual times. In a realistic situation, these times are rather given as time intervals corresponding to geological stages for fossil ages or to hypotheses for the origin of the diversification. Actually, we give the possibility for the user to provide only the lower bound for the origin of the diversification (in this case, the upper bound is given by the most ancient fossil age).

Let us first remark that it is possible to explicitly integrate the probability density of patterns of types **a**, **b** and **c** over all starting times *s* between two given times *u* and *v*. Namely, we have that

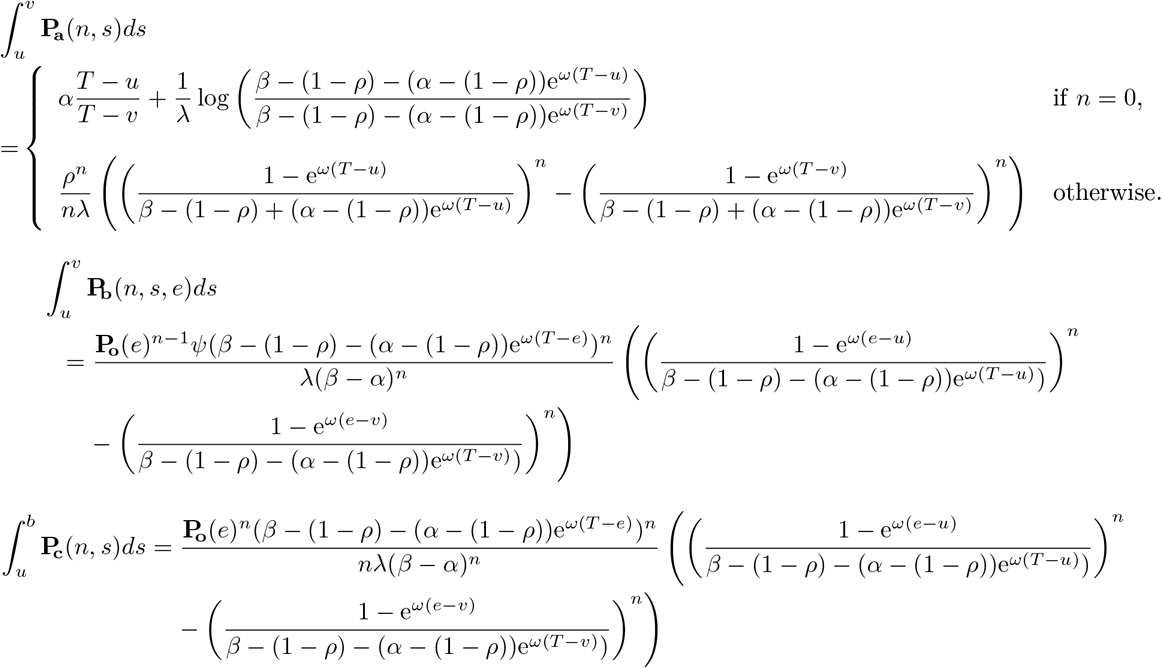

Dealing with uncertainty on the origin of diversification (i.e., the base of the root branch of the tree) is performed by replacing the probability density of the pattern starting at the origin by its integral between the two bounds *u* and *v* provided by the user or between *u* and the most ancient sampled fossil age if no upper bound is provided for the origin of diversification.

Since integrating all the possible fossil ages over the corresponding geological ranges is more complicated even when assuming an uniform distribution of the ages over their ranges (we plan to investigate this possibility soon), we have to sample them in order to make the divergence time distribution take into account uncertainty of the fossil ages expressed as ranges. In order to seed up the convergence of the distribution, we devised an importance sampling scheme based on the Metropolis-Hasting algorithm described in Appendix D (in a standard uniform sampling scheme, most of the iterations are not very useful because some fossil age configurations lead to very low tree-fossil probabilities, and these influence little the divergence time distributions).

In an earlier version of this work, the divergence time distributions were computed with regard to the maximum likelihood estimates of the speciation, extinction and fossilization rates. The divergence times distribution are now obtained by integrating over all the possible values of these parameters by assuming improper uniform priors, still by using the importance sampling procedure presented in Appendix D.

## EMPIRICAL EXAMPLE

### Dataset Compilation

Our dataset represents the fossil record of Cotylosauria (the smallest clade that includes Amniota and their sister-group, Diadectomorpha) from their origin (oldest record in the Late Carboniferous) to the end of the Roadian, which is the earliest stage of the Middle Permian. It represents the complete dataset from which the example presented in Didier *et al.* (2017) was extracted. Because of computation speed issues that we have now overcome, Didier *et al.* (2017) presented only the data on Eupelycosauria, a subset of Synapsida, which is one of the two main groups of amniotes (along with Sauropsida). Thus, the dataset presented here includes more taxa (109 taxa, instead of 50). We also incorporated the ghost lineages that must have been present into the analysis. However, in many cases, the exact number of ghost lineages that must have been present is unclear because a clade that appears in the fossil record slightly after the end of the studied period (here, after the Roadian) may have been present, at the end of that period, as a single lineage, or by two or more lineages, depending on when its diversification occurred. This concerns, for instance, in the smallest varanopid clade that includes *Heleosaurus scholtzi* and *Anningia megalops* (four terminal taxa). Our computations consider all possible cases; in this example, the clade may have been represented, at the end of the Roadian, by one to four lineages.

The data matrix used to obtain the trees is a concatenation of the matrix from Benson (2012), the study that included the highest number of early synapsid taxa, and of that of Müller and Reisz (2006) for eureptiles. However, several taxa studied here were not in our concatenated matrix. We specified conservatively their relationships to other taxa using a skeletal constraint in PAUP 4.0 (Swofford 2003), as we reported previously (Didier *et al.* 2017). The skeletal constraint reflects the phylogeny of Romano and Nicosia (2015) and Romano *et al.* (2017) for Caseasauria, Spindler *et al.* (2018) for Varanopidae, Brink *et al.* (2015) for Sphenacodontidae, and Brocklehurst (2017) for Captorhinidae. Note that our tree incorporates only taxa whose affinities are reasonably well-constrained. Thus, some enigmatic taxa, such as *Datheosaurus* and *Callibrachion*, were excluded because the latest study focusing on them only concluded that they were probably basal caseasaurs (Spindler *et al.* 2016). The recently described *Gordodon* was placed after Lucas *et al.* (2018). The search was conducted using the heuristic tree-bisection-reconnection (TBR) search algorithm, with 50 random addition sequence replicates. Zero-length branches were not collapsed because our method requires dichotomic trees. Characters that form morphoclines were ordered, given that simulations and theoretical considerations suggest that this gives better results than not ordering (Rineau *et al.* 2015), even if minor ordering errors are made (Rineau *et al.* 2018). The analysis yielded 100 000 equally parsimonious trees; there were no doubt more trees, but because of memory limitation, we had to limit the search at that number of trees. Benson (2012) had likewise found several (more than 15 000 000) equally parsimonious trees, and we had to add additional taxa whose relationships were only partly specified by a constraint, so we logically expect that our dataset would yield more trees. We performed our analyses on a random sample of 1000 equiparsimonious trees (one of these trees is displayed in Figure 6). As in our previous study (Didier *et al.* 2017), these trees do not necessarily represent the best estimate that could possibly be obtained of early amniote phylogeny if a new matrix were compiled, but that task would represent several years of work (Laurin and Piñeiro 2018) and is beyond the scope of the current study. We have placed parareptiles and varanopids in their traditional position, even though doubts about this have been raised by recent analyses (e.g., Laurin and Piñeiro 2018; Ford and Benson 2020). The analyses performed here can be repeated in the future as our understanding of amniote phylogeny progresses. In our dating analyses, all taxa were considered to represent tips (no fossils were placed on internal branches). It is conceivable that a few of the fossils included in our dataset represent actual ancestors, but the sensitivity analyses carried out by Didier *et al.* (2017) suggest that this should have a negligible impact on our results. The data are available in the supplementary information.

**Figure 6:**
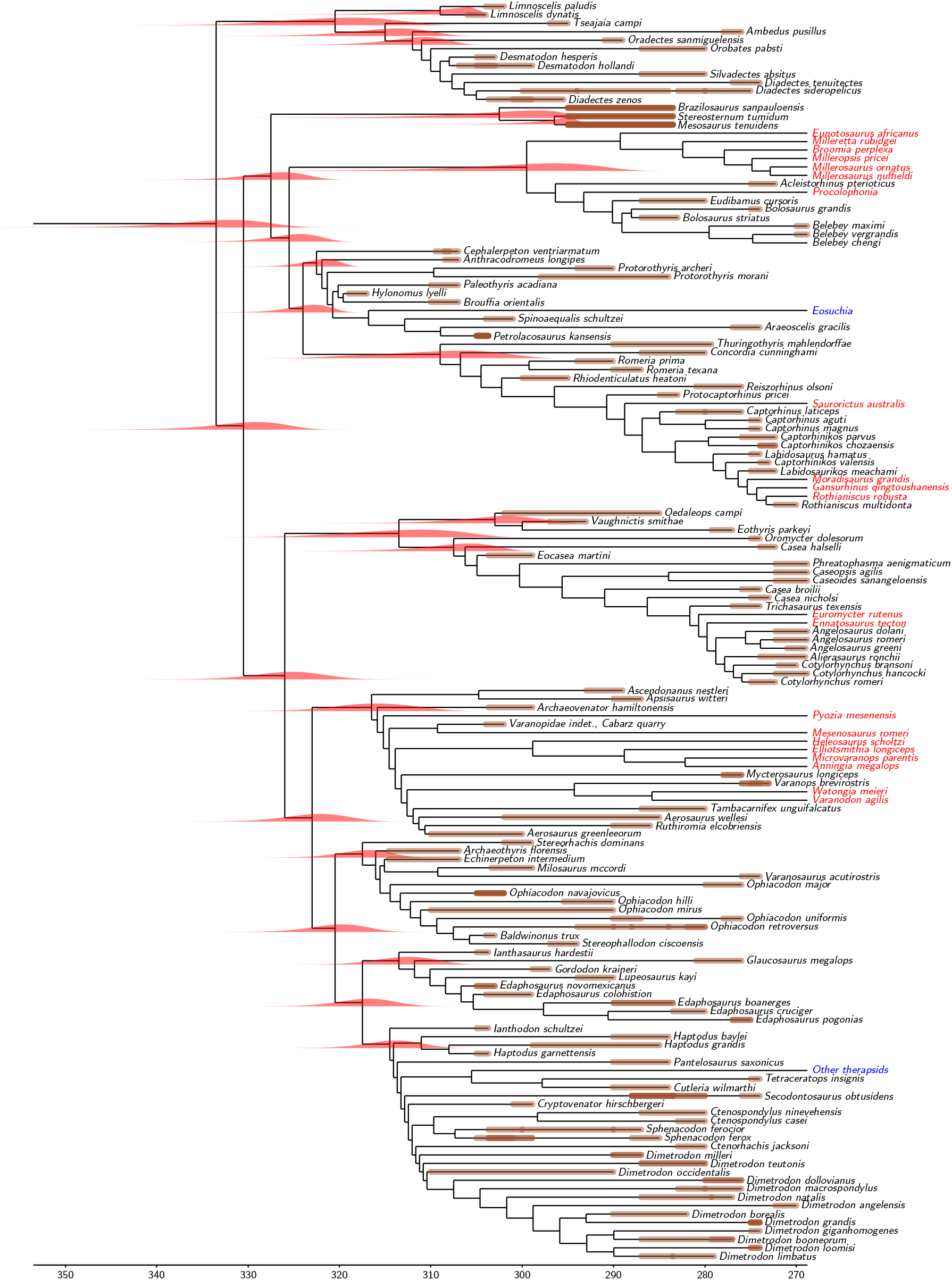
One of the 100 equally parsimonious trees used in our analyses with some divergence time distributions displayed at the corresponding nodes. The distributions shown here (contrary to those shown in Figures 7 and 8) were computed from this tree only (i.e., without taking into account the 999 other equiparsimonious trees). The brown highlighting of the branches represents the bounds of the range of plausible ages of fossil occurrences; darker brown indicates more than one record in the same time interval. Some taxa, displayed in red for extinct ones and in blue otherwise, lack a fossil record in the sample period (which ends at the end of the Roadian, at 268.8 Ma) but are included if their affinities suggest that a ghost lineage must have been present.

### Dealing with Fossil, Root Age, and Tree Topology Uncertainty When Estimating Nodal Ages

As in Didier *et al.* (2017), we used a flat distribution between upper and lower bounds on the estimate of the age of each fossil. That age was assessed by looking at the stratigraphic position of the fossiliferous localities in the primary literature. That was converted into absolute ages using the latest available geological timescale (Ogg *et al.* 2016), if absolute age data was not given directly in the source papers, or if the scale had been updated since then. For the taxa that were present in the analysis presented in Didier *et al.* (2017), the boundaries of the range of stratigraphic ages were not modified. However, our new analyses are based on a more inclusive set of taxa.

Our method also requires inserting a prior on the origin of the diversification, i.e., on the starting time of the branch leading to the root (here, Cotylosauria). We set only the lower bound of this origin and assume a flat distribution between this origin and the most ancient (sampled) fossil age (here that of *Hylonomus lyelli* between 319 and 317 Ma), a fairly basal eureptile sauropsid (Müller and Reisz 2006; Matzke and Irmis 2018). To study the robustness of our estimates to errors in this prior, we repeated the analysis with several origins that encompass the range of plausible time intervals and beyond. Recent work suggests that the Joggins Formation, in which the oldest undoubted amniote (*Hylonomus*) has been recovered (Carroll 1964; Davies *et al.* 2006; Falcon-Lang *et al.* 2006), is coeval with the early Langsettian in the Western European sequence, which is about mid-Bashkirian (Carpenter *et al.* 2015), around 317-319 Ma (Utting *et al.* 2010; Raine *et al.* 2015). In fact, recent work suggests more precise dates of between about 318.2 and 318.5 Ma (Utting *et al.* 2010; Rygel *et al.* 2015), but we have been more conservative in putting broader limits for the age of this formation, given the uncertainties involved in dating fossiliferous rocks. Thus, we have set the lower bound of the origin of diversification to 330 Ma, 340 Ma, 350 Ma, 400 Ma and 1 000 Ma to assess the sensitivity of our results to the older bound or the width of the interval. The lower bound of 1 000 Ma is of course much older than any plausible value, but its inclusion in our analysis serves as a test of the effect of setting unrealistically old lower bounds for the root age.

We deal with the uncertainty on the phylogenetic tree topology by averaging over the 1000 equiparsimonious trees provided in the dataset. The distribution displayed below were obtained by using the importance sampling procedure presented in Appendix D.

### Results

Figure 7 displays the posterior distribution histograms of the speciation, extinction and fossilization rates. The maximum likelihood estimates computed in Didier *et al.* (2017) from an earlier version of the dataset are consistent with the posterior distribution obtained here, rather in their lower parts. These distribution suggest a speciation rate slightly higher than the extinction rate.

**Figure 7:**
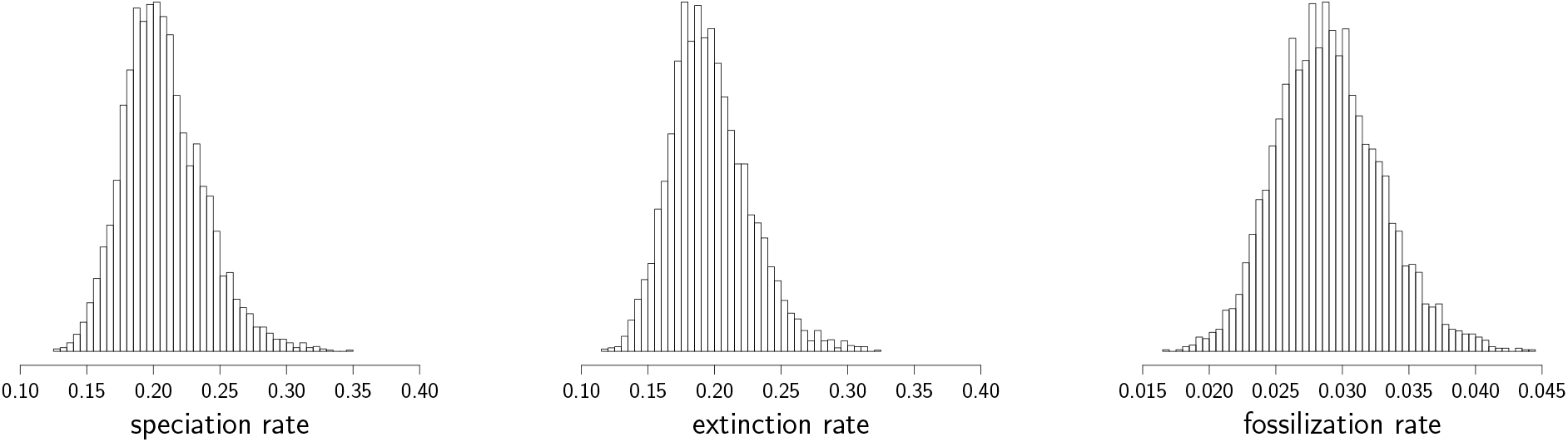
Posterior distribution histograms of the speciation, extinction and fossilization rates (events per lineage and per Ma).

Though the exact divergence time distributions can be directly computed in a reasonable time, it may be useful to fit them with standard distributions, for instance in order to use it in molecular dating software which do not implement their exact computation. Figure 8 shows that the divergence time distributions with lower bounds of the origin of diversification set at 1 000 to 350 Ma can be reasonably approximated by shifted reverse Gamma distributions with shape parameter *α*, scale parameter *θ* and location parameter *δ*, i.e., with the density function:

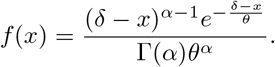

**Figure 8:**
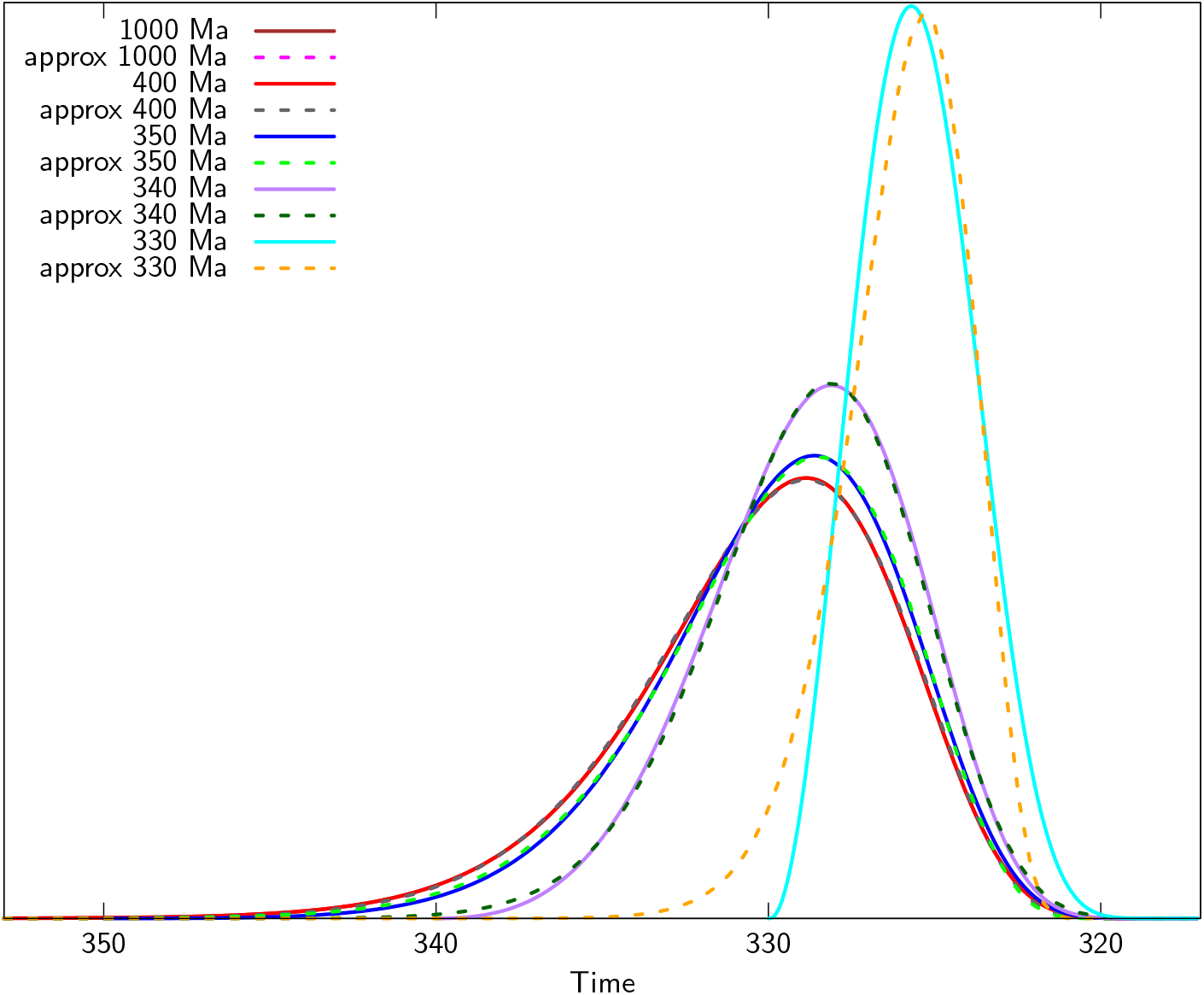
Probability density of the Amniota divergence time for several lower bounds of the age of origin of diversification averaged over all the possible fossil ages and speciation, extinction and fossilization rates with (improper for the rates) uniform priors (plots of the probability densities for 1 000 and 400 Ma overlap so tightly that the former is not visible).

Table 1 displays the best fitting parameters of the shifted reverse Gamma distributions plotted in Figure 8.

**Table 1:** Best fitting parameters of the shifted reverse Gamma distributions approximating the Amniota divergence time distributions displayed in Figure 8.

| Lower bound | $\alpha$ | $\theta$ | $\delta$ | Mean SSR |
| --- | --- | --- | --- | --- |
| 1000 | 8.377 | 1.377 | -318.702 | $1.321 \times 10^{-7}$ |
| 400 | 8.377 | 1.378 | -318.702 | $1.320 \times 10^{-7}$ |
| 350 | 7.501 | 1.392 | -319.463 | $8.167 \times 10^{-7}$ |
| 340 | 24.831 | 0.633 | -313.057 | $1.295 \times 10^{-6}$ |
| 330 | 8.500 | 0.662 | -320.314 | $2.260 \times 10^{-4}$ |

Note that shifted reverse Gamma distributions do not always approximate correctly divergence time distributions. This is in particular the case for the distributions obtained from the lower bounds 330 and 340 of the time origin, but also for those associated to several nodes in Figure 6.

#### Age estimates

Our results show remarkable robustness to variations in root age prior specification when the lower bound of the origin of diversification is far enough to the most ancient fossil age (Fig.8). The probability distributions of the age of Amniota obtained with origins 1 000 or 400 Ma are so close to each other that the curves are superimposed over all their course and thus, only one of these two curves is visible. Assuming that the lower bound is 350, 340 or 330 Ma predictably yields slightly narrower distributions, but the peak density is at a barely more recent age (around 328 to 330 Ma). Whatever the time origin, the probability density of the age of Amniota always dwindles from its peak to near 0 before reaching the age of 347 Ma. More importantly, the curves show that when the specified time of origin is at least 350 Ma, the probability density falls to near 0 well before reaching the time of origin, which suggests that the latter does not strongly constrain the result. This is even more obvious when looking at the peak density, which shifts very little (about 1-2 Ma) between times of origin of 340, 350, 400, and 1000 Ma. All this suggests that Amniota probably appeared approximately in the middle of the Carboniferous, which is fairly congruent with the fossil record, given that it indicates a minimal age around 318-319 Ma, only about 10-12 Ma less than the peak density.

## DISCUSSION

Our estimates of the rates of cladogenesis (speciation), extinction and diversification are more than 50% higher than to those obtained for a subset of our data (50 taxa out of the 109 used here) in Didier *et al.* (2017), and the fossilization rates are slightly lower. These moderate differences are not surprising given that we have expanded the taxonomic sample and made minor modifications to the method. This difference may reflect local variations in turnover rate in the tree, something that our method cannot yet test; but clearly progress could be made by estimating this variability in subsequent developments.

The method provided here is, to our knowledge, the first one able to compute the exact distributions of divergence times from fossil ages and diversification and fossilization rates. Previous approaches only allowed to sample these distributions by using Monte Carlo Markov Chain approaches. Our computation is fast, with a time complexity cubic with the size of the phylogenetic tree, and can handle hundreds of taxa. Divergence time distributions obtained from fossil ages through methods such as ours are natural choices to calibrate evolutionary models of molecular data and to be used as priors for phylogenetic inferences.

Our method is also useful to time-scale paleontological trees. Tip dating has been increasingly used for this (e.g., Ronquist *et al.* 2012a; Dembo *et al.* 2015; Lee and Yates 2018), but the recent demonstration (Goloboff *et al.* 2018) that most phenotypic characters fit poorly the Markov model of evolution that is currently used in tip dating raises doubts about the reliability of this approach and highlights the interest in developing additional approaches, such as using birth-and-death models. Development of such methods is timely because much progress has been made in the last decades in understanding the phylogeny of extinct taxa, as shown by the growing number of paleontological papers that included relevant phylogenetic analyses (e.g., Romano and Nicosia 2015; Bardin *et al.* 2017; Brocklehurst 2017). In turn, these progresses have triggered exciting developments of various methods to better time-scale paleontological trees (Hopkins *et al.* 2018). The methods mentioned above are only the latest in a long series of developments in this field. Notable earlier studies in this field include Hedman (2010), who provided a simple method (which does not use birth-death processes) to constrain clade age using a series of stratigraphically older sister-taxa.

Scaling paleontological trees has become important because such timetrees (and timetrees incorporating both extant and extinct taxa) have increasingly been used to tackle various evolutionary problems, such as evolution of biodiversity and evolutionary model (e.g., Ascarrunz *et al.* 2019) and evolutionary rate of phenotypic characters (Ascarrunz *et al.* 2016; Zhang and Wang 2019). The simulations of Bapst (2014) have shown that estimated rates of character change can be strongly over-estimated when tree scaling is wrong.

Our analyses suggest that the fossil record of early amniotes may not be as incomplete as previously feared. This is despite the fact that the fossil record of continental vertebrates is relatively poor in the Early Carboniferous (in “Romer’s gap”) a bit before the first amniote fossil occurrence (Romer 1956; Coates and Clack 1995; Marjanović and Laurin 2013). Thus, there was a possibility that amniotes had a much older origin and an extended unrecorded early history in Romer’s gap. This had led Müller and Reisz (2005) to argue that the appearance of amniota was too poorly documented in the fossil record to be useful as a calibration constraint for molecular dating studies. However, our results suggest that these fears were exagerated; the fossil record of amniotes appears to start reasonably soon after their origin, with a gap of no more than about 20 to 25 Ma separating amniote origins from the first recorded fossil occurrence. A caveat preventing more definitive conclusions on this point is that our analyses assume constant rates throughout the tree and throughout the studied period. Thus, if the fossil record were poorer in the early parts of the interval (e.g., in the Late Carboniferous) or if diversification accelerated over time, our results might be biased. Further developments to accommodate variations in rates through time are in progress and should improve the robustness of our estimates.

Our results about the age of Amniota should prove useful for a wide range of node-based molecular dating studies that can incorporate this calibration constraint. The main objection by Müller and Reisz (2005: 1074) against use of the age of Amniota in molecular dating (the uncertainty about the maximal age of the taxon) has thus been lifted; we now have a reasonably robust probability distribution that can be used as prior in node dating. Indeed, our probability distributions for the age of Amniota probably make it one of the best-documented calibration constraint so far. The probability distributions are fairly well-constrained (with a narrower distribution than many molecular divergence age estimates) and show surprisingly little sensitivity to the maximal root age prior, which is reassuring. A 95% probability density interval of Amniota, using a 350 Ma maximal root age constraint, encompasses an interval of about 20 Ma. By comparison, a 95% credibility interval of nodes of similar ages, such as Lissamphibia, in Hugall *et al.* (2007: table 3) encompasses 38 Ma or 56 Ma, depending on whether these are evaluated using the nucleotide or aminoacid dataset. Some nodes in Hugall *et al.* (2007: table 3) are better constrained, probably because they are closer to a dating constraint. Thus, Hugall et al. report a 95% credibility interval for Tetrapoda that encompasses a range of 24 Ma and 32 Ma, for the nucleotide or aminoacid datasets. This is only moderately broader than our 95% interval, but the width of the intervals reported in Hugall et al. (2007: table 3) are underestimated because they reflect a punctual estimate (at 315 Ma) of the age of Amniota, which is used as the single calibration point. Pyron (2011: table 1) reports 95% intervals of 54 Ma for the age of Lissamphibia using Total Evidence (tip) dating. Similarly, Ronquist *et al.* (2012a: fig. 9) obtained 95% credibility intervals of about 50 Ma for major clades of Hymenoptera using tip (total evidence) dating, and substantially broader intervals using node dating. Recently, Ford and Benson (2020) estimated the age of Amniota at about 324.5 Ma with a 95% credibility interval ranging from about 316 to 328 Ma using a method incorporating a morphological clock (tip dating) and the FBD. These results are broadly congruent with ours to the extent that the credibility intervals overlap widely. Thus, we believe that our estimates are fairly precise, when comparisons are made with molecular estimates that consider a similarly broad range of sources of uncertainty. However, such comparisons are hampered by the fact that phylogenetic uncertainty is handled differently by various methods (directly, by integrating fossils into the phylogenetic analysis in tip dating and here, but only indirectly in node dating, by relying on the oldest undisputed fossil of each clade), and by the fact that each method has a different mix of strengths and weaknesses. Among the weaknesses, our method assumes that fossilization and diversification are constant over time; tip dating assumes that phenotypic characters evolve in a clock-like manner and can be modeled adequately by the Mk model (an assumption that was shown to be unreasonable by Goloboff *et al.* (2018); node dating discards much fossil data and requires inputting node age priors about which we have little information, except for the minimal age. Also, in empirical studies, given that we may never know the actual divergence times, it is impossible to know which methods give the most accurate results. Simulations would be helpful to study this question, but they are well beyond the scope of this study.

**Figure 9:**
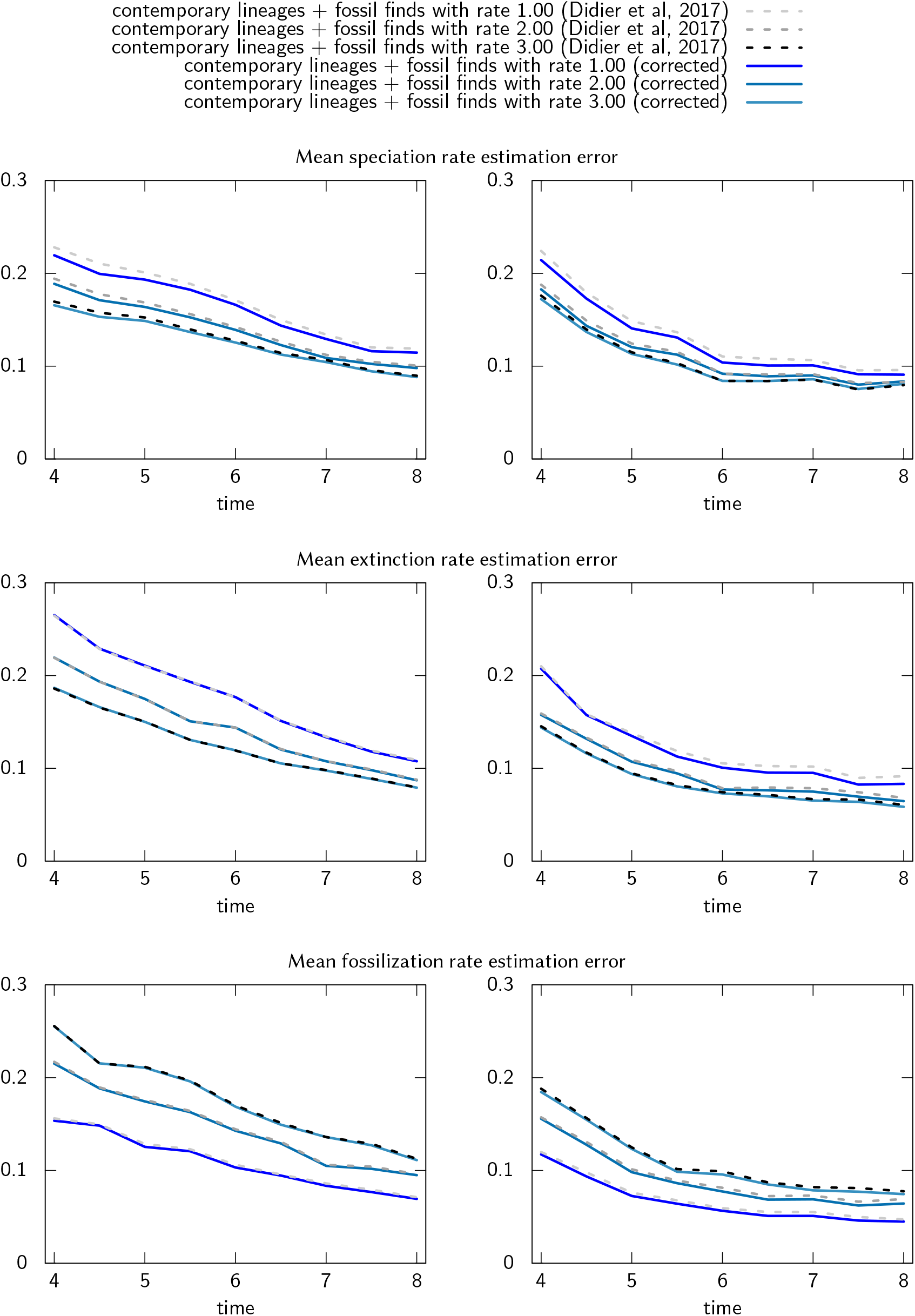
Mean absolute error of speciation (cladogenesis), extinction and fossil discovery rate estimation *versus* the simulation (evolutionary) time over 1 000 simulated trees with 1.5/1 (left) and 2.0/1 (right) speciation/extinction rates and fossil discovery rates from 1 to 3.

Our estimates also suggest that the way in which this constraint (Amniota) was used in most molecular dating studies was not optimal. Indeed, most (e.g., Hedges *et al.* 1996; Hugall *et al.* 2007) have set a prior for this node centered around 310 to 315 Ma, whereas our analyses suggest that the probability peak is approximately around 330-335 Ma. It will be interesting to see how much change will be generated by these new data, and with similar data on other calibration constraints that can be obtained with our new method.

Our method can be applied to any clade that has a good fossil record and a sufficiently complex phenotype to allow reasonably reliable phylogenetic analyses to be performed. In addition to vertebrates, this includes, minimally, many other metazoan taxa among arthropods (Ronquist *et al.* 2012b), echinoderms (Sumrall 1997) and mollusks (Merle *et al.* 2011; Bardin *et al.* 2017), among others, as well as embryophytes (Corvez *et al.* 2012). With new calibration constraints in these taxa (and possibly others), the timing of diversification of much of the eukaryotic Tree of Life should be much better documented.

## ACKNOWLEDGMENTS

We thank Michael C. Rygel (SUNY Potsdam) for sending papers about the stratigraphy of the various formation represented in the Joggins locality. P. Drapeau and M. Fau helped to compile the data for the empirical example.

## A TABLE OF THE NOTATIONS

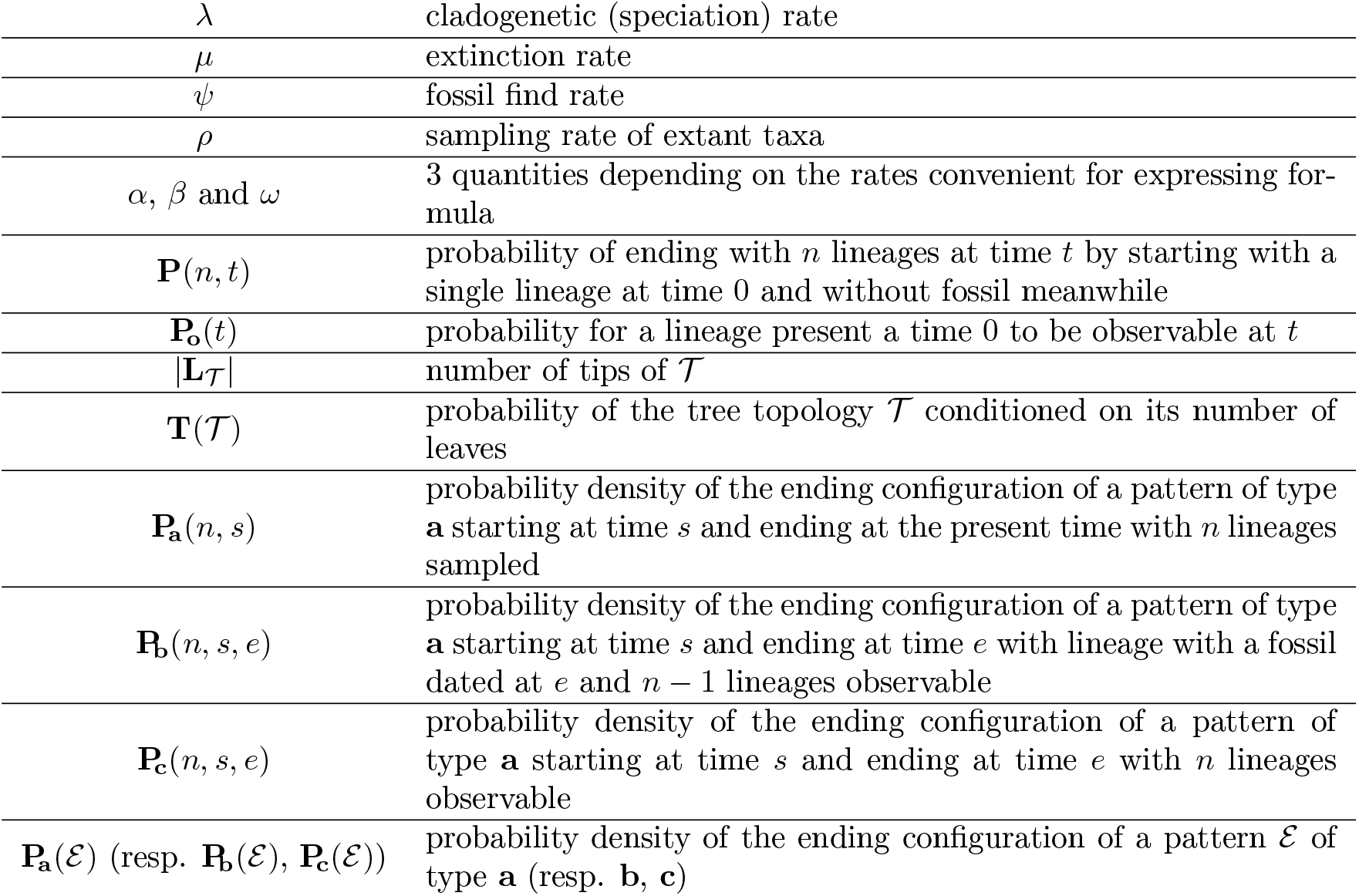

## B EQUIVALENCE BETWEEN TWO TREE TOPOLOGY DISTRIBUTIONS

In Didier *et al.* (2017), the probability 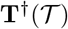 of a tree topology 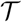 arising from a lineage homogeneous process (conditioned on its number of tips) was expressed as

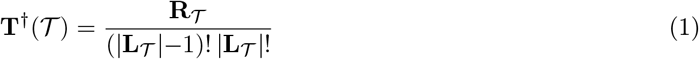

where 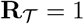 if 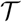 is made of a single lineage and, putting *a* and *b* for the two direct descendants of the root of 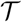,

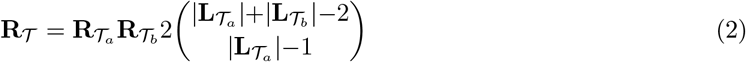

otherwise.

If 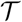 is made of a single lineage, both the probability 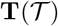 from Harding (1971) and 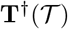 are equal to 1.

Otherwise, by substituting Equation 2 in Equation 1, we get that

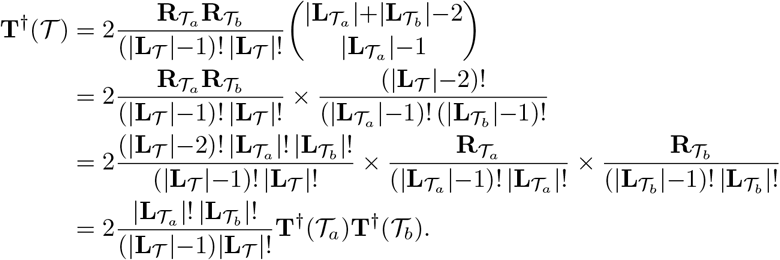

It follows that the probability 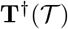 of Didier *et al.* (2017) can be expressed with the same recursive formula as the probability 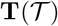 of Harding (1971), recalled in Theorem 1. Since moreover the recursive computations of the probabilities of Harding (1971) and Didier *et al.* (2017) have the same initial condition (i.e., that of trees made of a single lineage), they are equal for all topologies 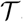. It is worth noting that the way in which these distributions have been derived is quite different. Arguments of Harding (1971) mainly rely on the labelling of the tree while those of Didier *et al.* (2017) are essentially based on the relative order of its divergence times.

## C IMPACT OF THE CORRECTION OF THE PROBABILITY DENSITY ON THE RATES ESTIMATION

In order to assess the impact of the error in the probability density of phylogenetic tree with fossils on the diversification rate estimation, we simulated trees and fossils following an FBD model with given rates and determine their maximum likelihood estimates on the simulations by using the computation from Didier *et al.* (2017) and the corrected one presented here. Thanks to the improvement of the complexity of the new algorithm, which we also adapted for re-implementing our former computation, we relaxed the filtering of the simulated trees. Namely, we now rejected trees with fewer than 10 extant taxa or with more than 5 000 clades, against 20 extant taxa and 1 000 clades in Didier *et al.* (2017). Unlike for simulations of Didier *et al.* (2017), we did not filter the simulated trees with regard to their complexity level (i.e., their expected computational time with the former algorithm).

Figure 9 displays the mean absolute error of speciation, extinction and fossil discovery rates obtained from the corrected method provided in this work compared to the former one of Didier *et al.* (2017), in a same set of simulated phylogenetic trees with fossils. Though we can observe a slight improvement for the estimates of the corrected computation, the accuracy of the estimates is essentially the same for the two methods.

## D A METROPOLIS-HASTINGS IMPORTANCE SAMPLING PROCEDURE

Our computation of the probability density of a phylogenetic tree topology with fossils with or without time constraints on its divergence times requires that both the topology and the fossil ages are exactly known, a situation which is not met in practice. Let us show how to deal with the case of our dataset, i.e., where a set Ω of equiprobable topologies are provided and where age of each fossil *f* is only known from its geological age interval I_*f*_. For all trees 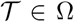 and all vectors of fossil ages **f** ∈ **I**, we have a phylogenetic tree topology with fossils 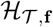 for which we are able to compute the probability density. All the topologies in Ω are assumed equiprobable. We make the additional assumption that the distribution of fossil ages is uniform on the product-set of the time intervals **I** = ⊗_*f*_ **I**_*f*_. Taking into account the uncertainty of the dataset, the divergence time distribution associated with a given internal node *n* is

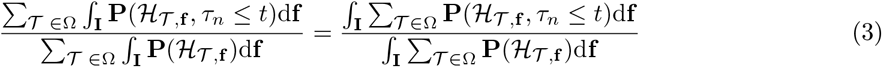

The quantities involved in the ratio above cannot be directly computed and have to be estimated, for instance by uniformly sampling the fossil ages in **I**. Being given *N* fossil age vectors **f**_1_, …, **f**_*N*_ uniformly sampled from **I**, the estimated divergence time distribution associated to *n* is

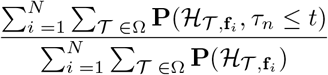

Unfortunately, because of its variance, the estimator above requires millions of samples to converge in our dataset. A usual way to tackle this issue is to perform importance sampling (see, e.g., Robert and Casella 2013). The general idea is to estimate the integral 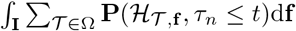 by sampling the fossil ages not uniformly but following a density q^∗^ biased toward the ages corresponding to high values of 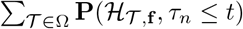. Since

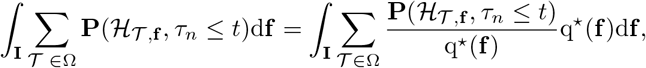

if 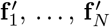 are independently sampled from density q^∗^, an estimator of this integral is

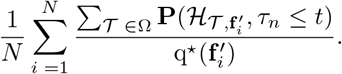

The fact that 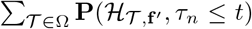 is bounded by (and strongly related to) 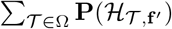 suggests that considering a density q^∗^(**f**) proportional to 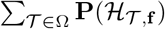 is suited for the importance sampling of 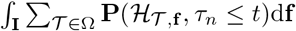. Let us set

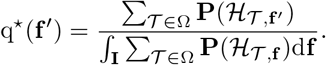

If 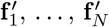 are iid samples with the density q^∗^ just above, the ratio of Equation 3 is estimated by

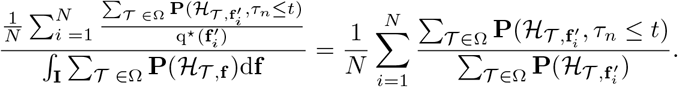

Since it is not possible to directly sample following the density q^∗^, which is known only up to the intractable normalization constant 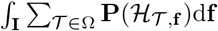, we used the Metropolis-Hastings algorithm (e.g., Robert and Casella 2013) to build a Markov chain with target density q^∗^. Given the *i*^th^ vector of fossil ages 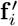 of this chain, we generate a candidate vector 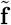 first by uniformly selecting a fossil *f* among all the fossils of the dataset and by replacing the corresponding entry of 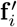 by a uniform sample in the interval centered on this entry of width equal to *α* time the with of the interval I_*f*_ that represents the uncertainty on the age of fossil *f* (i.e., we use a usual sliding window proposal with reflection; *α* is a user-defined parameter). This proposal transition kernel is basically symmetrical, i.e., getting the candidate vector 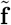 from 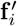 has the same probability as getting the candidate vector 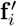 from 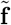. The next fossil age vector 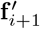 in the Markov chain is

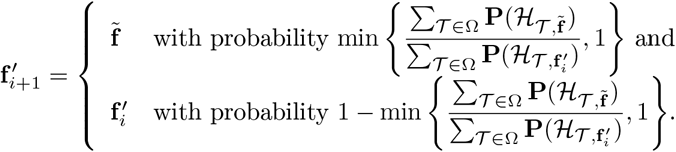

In the case where the speciation, extinction and fossilization rates of the model are not given *a priori* (e.g., estimated by maximum likelihood), we sum over all their possible values by assuming that they all follow the improper uniform distribution over [0, +℞] and by following a similar importance sampling procedure. In plain English, the importance sampling is now perform over both the fossil age vector **f** and the model rates Θ = (*λ, μ, ψ*) by using the biased density q^∗^(**f**, Θ) proportional to 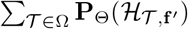, where 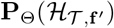 is the probability of the phylogeny with fossils 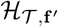 under the FBD model with rates Θ = (*λ, μ, ψ*).

The proposal mechanism of the Markov chain used to sample according to this density first draws a Bernoulli random variable with a probability *β* (provided by the user) according to which either a change in a fossil age or a change in the speciation, extinction or fossilization rate is proposed. The change in a fossil age is proposed in the exact same way as just above. The change in a rate is proposed first by selected randomly the speciation, extinction or fossilization rate according to user-defined probabilities then by using a sliding window of fixed length centered on the current selected rate.

The densities presented in the results section were obtained from the MCMC procedure presented just above by discarding the first 10000 iterations then by keeping only 1 iteration over 300. We consider a window of width equal to 80% of the length of the age interval I_*f*_ to propose a change in the age of fossil *f* and windows of size 0.1, 0.1 and 0.01 to propose changes in the speciation, extinction and fossilization rates respectively. We propose a change in a fossil age with probability 0.75 and a change in the speciation, extinction or fossilization rate otherwise (the rate to be changed among the the speciation, extinction and fossilization rate is then drawn uniformly). We run the MCMC procedure until get 2000 iterations. The iterations thus obtained pass all the convergence tests of the coda R package with an effective size greater than 200 for all the parameters (fossil ages and rates, Plummer *et al.* 2006). All the parameters of the MCMC can be modified by calling the corresponding R function in the package.

